# The structural quality of maternal health services in primary health care facilities in Tanzania: Findings from a baseline study

**DOI:** 10.1101/699553

**Authors:** Ntuli A. Kapologwe, Albino Kalolo, Naomi H. Isanzu, Josephine Borghi, Stephen M. Kibusi

**Author notes:** Corresponding author details and complete co-author email addresses, Ntuli A. Kapologwe. Josephine Borghi -, Albino Kalolo -, Stephen Kibusi.

## Abstract

**Background:** Structural quality of maternal health services remains a key indicator of highly performing health care system. Evidence attest to the fact that introduction of the new interventions in the health care system does not necessarily lead into improvement of the target outcome, such as quality of health services delivered. This study aimed at assessing the structural quality of maternal health services prior to introduction of Direct Health Facility Financing (DHFF) program.

**Methods:** This was a cross-sectional study, conducted in 42 public primary health facilities between January and mid February 2018. Observational were used to collect the data from health facilities. The collected information was on privacy, hygiene and sanitation, obstetric emergences, sterilization, maternal death audit reviews and waste management. Collected data were analyzed by using SPSS.

**Results:** All 42 (100%) primary health facilities that were assessed were public primary health facilities, of which 14 (33.3%) were health centers and 28 were dispensaries. The furthest primary health facilities from the district head office were 140 Kms and the nearest 2 Kms. Focusing on; - privacy, hygiene and sanitation, obstetric emergences, sterilization, maternal death audit reviews and waste management assessed eight areas of Structural qualities. Majority (68.9%) of Health Centers has less than 39 skilled staff while some of them they have up to 129 health service providers and majority (92.8%) of Dispensaries have less than 15 staff and some have 1 staff.

By comparing Dispensary and Health center performances on structural quality indicated relatively low differences among the attributes assessed. Specifically, they did not show statistical significant differences except for obstetric emergencies (p < .005), sterilization (p=. 034) and overall structural quality (p=. 018). With regard to rural-urban performance on structural quality, there was no statistical significant difference on total performance. Similarly, there was no significant differences between rural and urban health facilities on other assessed attributes of structural quality (p >.05) except for sterilization in which urban facilities performed significantly higher than the rural facilities [M=41.2, SD=27.7, 61.3, SD=28.4, respectively (p= .028)]. On the other hand, marginal differences were observed on individual assessed attributes. For examples, rural facilities performed relatively higher than urban ones on privacy (41.2 and 32.0), maternal death reviews (31.4 and 30.7) and waste management (49.0 and 47.3) respectively.

**Conclusion:** Generally facilities performed low on the structural quality indicators of maternal health services provision however; they had high performance on sterilization and emergence obstetric care.

## Background

Quality of care can be defined as ‘degree to which health services for individuals and populations increase likelihood of desired health outcomes and consistent with current professional Knowledge, [1].

‘Structures’ refers to health system characteristics that affect the system’s ability to meet the healthcare needs of individual patients or a community. Structural indicators describe the type and amount of resources used by a health system or organization to deliver programs and services, and indicators relate to the presence or number of staff, clients, funds, beds, supplies, and buildings including those which offers safe surgery facilities. Primary health care is an important entry point into the healthcare system particularly in Tanzania; it is therefore important to invest in its quality structures so that to improve health service utilization of the population at the same reducing morbidities and mortalities [2]. Donabedian defines the quality of services by focusing mainly on three domains namely structure, process and outcome [3]. Body of evidence suggests the association existing between the structures and the better quality of health services being offered by those facilities [4,5]. So structural maternal health services are not exceptional to the fact.

Globally, it is estimated that 303,000 maternal deaths occurred in the year 2015, with sub Saharan Africa contributing up to 70% of maternal mortality worldwide [6]. Of all the Millennium Development Goals (MDG), the goal number five which aimed to reduce maternal mortality by ¾ by 2015, it is the one which is not yet to be achieved to date [30]. Tanzania ranks among the countries with the highest maternal mortality rates worldwide [7]. The Tanzanian estimated maternal mortality ratio is 556/100,000 live births [8]. Tanzania has introduced several strategies that aims at reducing the maternal mortality, some of them are;-Health Sector Strategic Plan-IV (2015-2020) and One Plan I & II [9–11]. Evidence shows that poor quality facility-based care is a major contributing factor to elevated maternal and neonatal morbidity and mortality rates [12].

Globally, most Structural quality improvement programs mainly focus on hospitals [13 – 17] and it is unclear if there are benefits and impacts that could be reproduced at the primary health care facilities (dispensary and health centre) levels. In order to change the status quo, Tanzania like any other Country has implemented different quality improvement strategies, such as maternal death audits, standard-based management and recognitions, all of which are geared towards improving maternal health outcomes [18 – 20]. Improvement of structural quality of maternal health at a health facility level is heavily dependent on resources availability such as health care workers, amenities, medicines, medical equipment and supplies and infrastructure [21].

In ensuring that the quality of maternal health services is maintained at all levels, the Government of Tanzania has developed quality improvement framework in health care (2011 – 2016) [22,23]. Furthermore, in 2014 the Government of Tanzania did introduce star rating program for the primary healthcare facilities so that to grade their performances and foster provision of quality maternal health services and other services to its general public, this was preceded by development of quality improvement plans at the individual primary healthcare facility level [24] which guided implementations of quality maternal interventions. In implementing the star rating strategy, all primary health care facilities are star rated from O star to 5 stars according to the set quantity and quality indicators of service provision. According to the first star assessment exercise which was done in 2014 of 6,272 health facilities which were assessed 2,263 (36.06%) had 0 star, 3,277 (51.64%) had 1 star, 690 (10.99%) had 2 stars, 77 (1.23%) had 3 stars, 5 (0.08%) had 3 stars and none had 5 stars [24]. There have been some other efforts to ensure that the quality of services at the primary health facility is maintained in the efforts like supportive supervision and clinical services audit [25]. The reason of studying maternal health-related indicators lies on the fact that, Tanzania is one of seven countries that accounted for 3 – 5% of global maternal deaths reported in 2010 [26].

This study is part of the larger before and after evaluation study [28] that aims to establish the effects of the Direct Health Facility Financing (DHFF) reforms and its pillars (Figure 1) on structural quality of maternal health. More specifically this study aimed to determine the status of structural quality of maternal health in primary health facilities prior to the implementation of the Direct Health Facility Financing (DHFF) program.

**Figure 1.**
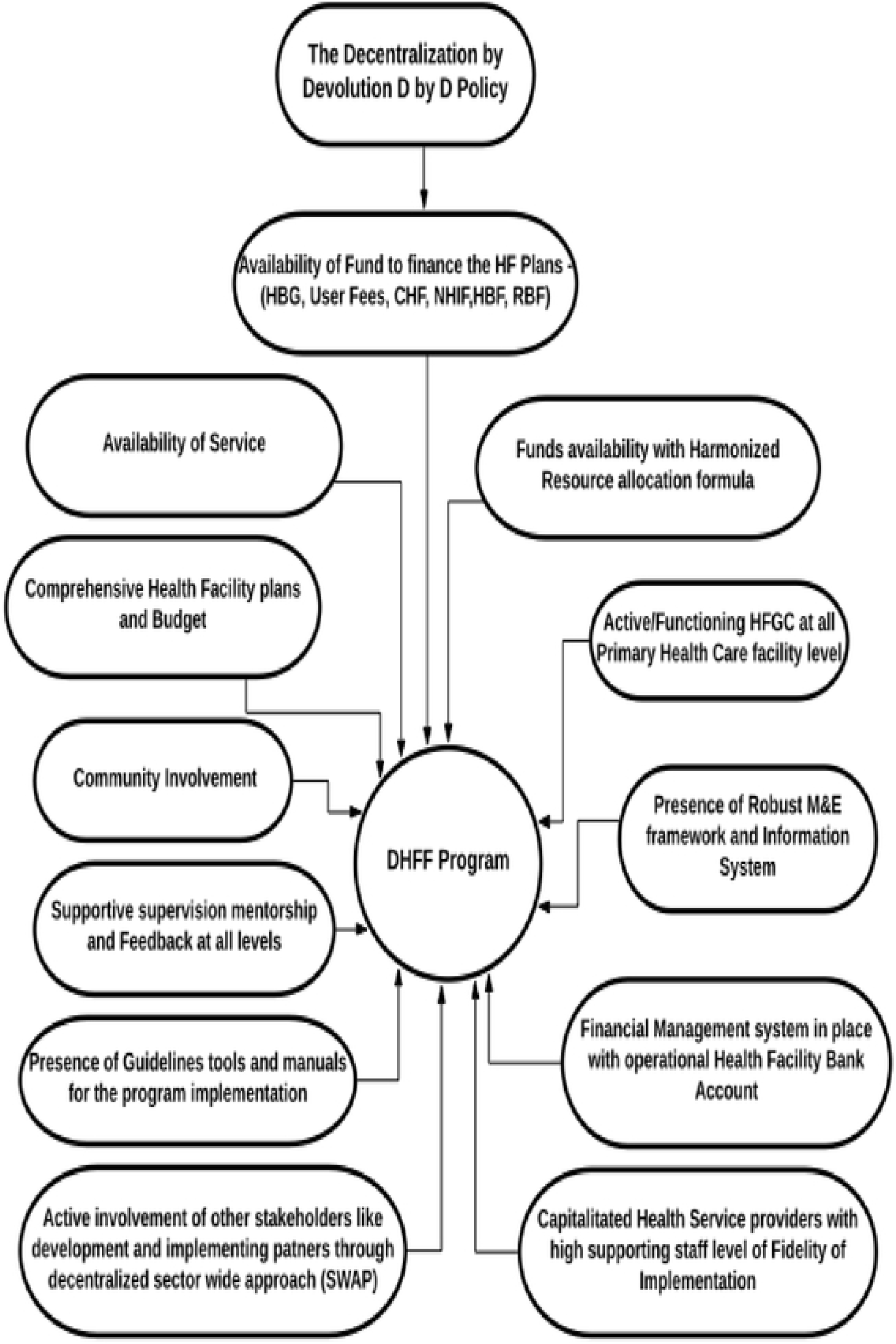
Pillars of the Direct Health Facility Financing

## Methods

### Settings

This study was conducted in fourteen (14) councils of seven (7) regions from seven (7) geographical zones. The reason for selecting 7 zones is to seek for country’s geographical representation. The similar approach has been done in major health studies in Tanzania [8]. This study was done in 3 primary public health care facilities (1 Health Center and 2 Dispensaries) in each of the 14 councils of Tanzania; a total of 42 primary health facilities will take part in this study.

This study is part of a larger study that aims at establishing the effects of DHFF on structural quality of maternal health services. Moreover, this study aims at establishing the baseline situation of the structural maternal health services of primary health facilities in Tanzania [28].

#### The organization of Health System in Tanzania

The health system of Tanzania is organized in a pyramid pattern. At the base of the pyramid there is the community followed by Dispensary and Health centres which constitute the primary health care. These are followed by District Hospital or Designated District Hospital that then followed by Regional Referral Hospitals, Zonal Hospitals, Specialized Hospitals and finally National Hospital. Meanwhile, there area total of 6,640 (53%) Dispensaries of which 4,554 (36%) are government owned. There are a total of 695 (15.7%) health centers of which 518 (11.6%) are government owned. The formal distinction between Dispensaries and Health Centers is that while Dispensaries exclusive provide out-patient care, a Health Center should be able to provide around the clock care to patients [29].

There are two types of primary health facility in Tanzania: dispensaries, providing a basic range of preventive, health promotion, curative and maternal and child health (MCH) care, and health centres, offering inpatient and a higher level of delivery care and staffed by a wider range of more qualified health workers up to 52 than in the dispensaries which have up to 20 health service providers [29]. The Alma Ata and the Astana Declarations further cemented the importance and evolution of PHC in 1978 and 2018 respectively [30,31], which emphasized the role of primary health care in service provision, especially PHC, to achieve an ambitious goal of health for all through quality, accessible and affordable health care services which does not render someone into catastrophic expenditure.

### Study design

This was a cross sectional study that aims at assessing the baseline situation of structural quality of maternal health services before DHFF program implementation [27]. The study was done between January and early February 2018.

### Sampling and sample size

Sampling was done using a four-stage sampling approach. The first stage included random selection of seven regions from the seven regions of Tanzania, clustered into seven geographical zones (each cluster constituted between 3 and 4 regions). In the second stage, selection of district councils was done and two district councils basing on stratification were selected into one urban and one rural, in their respective regions. The third stage comprised of selection of health facilities to be included into the study, they were selected at random from each strata of each district council in the 7 regions. A total of 3 primary health facilities were selected randomly from each district’s list of each type of Public Primary Health Care Facilities (PPHCF) (i.e. 1 Health Centers and 2 Dispensaries) (http://hfrportal.ehealth.go.tz) [32] i.e. Making a total of 42 health facilities (14 health centers and 28 dispensaries).

The sample size for health facilities was determined by taking a minimum sample size of 42. The study sample size calculation based on the purpose and nature of this study and the population under scrutiny was considered to be adequate. Thus, a minimum sample size of thirty (30) can be considered if some form of statistical analysis is to be done [33, 34].

For the health facilities to participate in the structural quality survey, the sampling frame were obtained with a list of all facilities in the district, from the sampling frame 42 HF were selected through multistage sampling technique and by stratification of health facilities into rural and urban. To each strata health facilities were randomly selected by using a table of random number.

### Data collection tools and procedures

The study measured the structural quality of maternal health services through observational checklist prior to the DHFF program inception and the Health Facility general information. The selection of these indicators based on the current challenges around maternal health services in Tanzania. The Observational Checklist is divided into seven parts namely Privacy, Hygiene and Sanitation, Labour ward, Obstetric emergencies, Waste management, Sterilization and Maternal deaths audit (Additional File 1). The database (data collection software) was developed to which all the data obtained from the study units were entered. This study used Mobile Data Collection (MDC). Observations data were captured on mobile phone entered into a pre-designed database.

### Observation checklist

A structured observation checklist adopted from that of Results Based Financing (RBF) [35] was used to collect data on structural quality of maternal health services. The RBF program is also implemented in more than 48 African Countries [36]. The information which were collected includes; facilitates which offers privacy, presence of hygiene and sanitation facilities, availability of appropriate equipment and material obstetric emergencies, availability of proper infrastructure and instruments for sterilization, facilities that completed maternal death audit reviews and have the action plan, and waste management is offered as per standard (additional File 1). In addition to that, the other information that was collected was the staffing level of individual primary health facility.

To ensure accuracy of the collected data and information, research assistants underwent four days training on data collection using a mobile device before taking part in pre-testing the tools. All selected facilities had GPS coordinates and all the data enumerators used tablets with GPS sensors. Mobile phone was used for data collection from the study areas. It was a web-based interface that allows real-time gathering of data and the principal investigator was monitoring the data collection exercise on daily basis to ensure for the integrity and quality of data. After the actual field survey collected data was then sent directly to the Gmail account app (which will acted as a server) after being filtered in the field.

## Data analysis

### Variables and measurements

Structural quality of maternal health services was determined by measuring mean of each of the 7 attributes of the structural quality indicators as stipulated in the observational checklist (Additional File 1).

The Structural Percent Score (SPS) for the seven attributes that had the minimum score which was less than the cut-off point for the acceptable performance and were regarded as *‘poor structural quality’* (Additional file 1). Those percent performances above the cut-off were categorized as *‘Good structural quality’*. The minimum SPS for each of the 7 attributes was calculated using the minimum structural quality scores (sqs) such as Privacy was determined by individual treatment /service delivery rooms that assessed based have full privacy during service provision had a total score of 4; Hygiene and Sanitation presence of clean and functioning disinfected toilet/s for patients, staffs and physically challenged people had a total score 7; Labor room focused on state of delivery room with essential equipment and supplies for quality service delivery and had a total of 14; Waste Management was assessed as per standard guidelines in clinical procedures rooms had a total score of 6; Maternal death audits in Primary Health Facilities that are completely and appropriately audited and action plan in place had a total score of 15; Obstetric emergences assessed based on availability of appropriate equipment and materials available to treat/manage patients with obstetric emergencies and had a total score 30; Sterilization assessment based on availability of proper sterilization of instruments and at total score of 3. All seven attributes had a total score of 79. SPS of the individual facility were multiplied by 100.0% and divided by 79 (Additional file 1).

All respondents scores for the seven attributes were converted to percent i.e. the scores divided by the maximum possible scores multiplied by 100%. These percent scores for the response categories were used as cut-off points to group the Structural quality performances of the primary healthcare facilities. As carried out in the similar study on structural aspect of quality of care [28, 37], individual elements were weighted based on RBF program on what is considered to be good or poor. On the guidance of RBF program operational manual and experiences from 7 regions implementing it and from other previous studies a standard of 60% was established to distinguish between health facilities providing good and poor structural quality [28,38, 39]. Points were then allocated to these characteristics, which in turn allowed calculation of an overall score for each of the attributes.

### Statistical analyses

The univariate and bivariate analyses were performed to describe the frequency, distribution and binary relationships. The unit of analysis was health facility. The data were visually inspected using the graphs for normality, linearity of relationships and the presence of outliers in the dependent variable and residuals. To check for the fulfillment of the first basic assumption i.e. normality of the dependent variable, Shapiro-Wilk W test for normal data was conducted and showed p ≥ 0.050 (p = 0.259).

Data was transferred into a Microsoft excel Database, and exported to Statistical package for Social Sciences (SPSS) version 21 for statistical analysis. Data cleaning was undertaken before actual statistical analyses.

The first step included a descriptive statistics (frequency, percentage, mean, and standard deviation) analysis of the structural quality of maternal health services. The observational check list that was used for the data collected was then used for guiding both descriptive and inferential statistics.

Multiple regression analysis was done to assess the relative effect of various explanatory variables or establish whether the predictor variables are independently associated with outcomes (dependent variable) of interest (Structural Quality attributes) was done to control/adjusting for any confounders as well.

### Ethics approval and consent to participate

Ethical approval for this study has been granted by the University of Dodoma (UDOM) ethical clearance committee and endorsed by National Institute of Medical Research (NIMR)-(Ref.No.NIMR/HQ/R.8a/Vol.IX/2740). Consent form was issued to participants prior to administration of the questionnaire.

## Results

### Health Facilities characteristics

A total of 42 (100%) primary health facilitates were assessed on structural quality maternal health services. Of all the facilities assessed 14 (33.3%) were health centers were as 28 (66.7%) were dispensaries. The furthest primary health facilities from the district head office were 140 kms and the nearest 2kms. Thirty three percent of health facilities were located in the rural areas (Table 1). Majority (68.9%) of Health Centers has less than 39 skilled staff while some of them they have up to 129 and majority (92.8%) of Dispensaries have less than 15 staff and some of them have 1 staff (Table 1).

### Structural Quality Maternal Health Services (sQMHS)

Shinyanga and Pwani regions had their structural quality performance 73.8% and 65.5% respectively as compared to other regions (Figure 2). Structural quality attributes that were assessed are: - facilities which offers privacy, presence of hygiene and sanitation facilities, availability of appropriate equipment and material obstetric emergencies, availability of proper infrastructure and instruments for sterilization, facilities that completed maternal death audit reviews and have the action plan, and waste management is offered as per standard.

**Figure 2:**
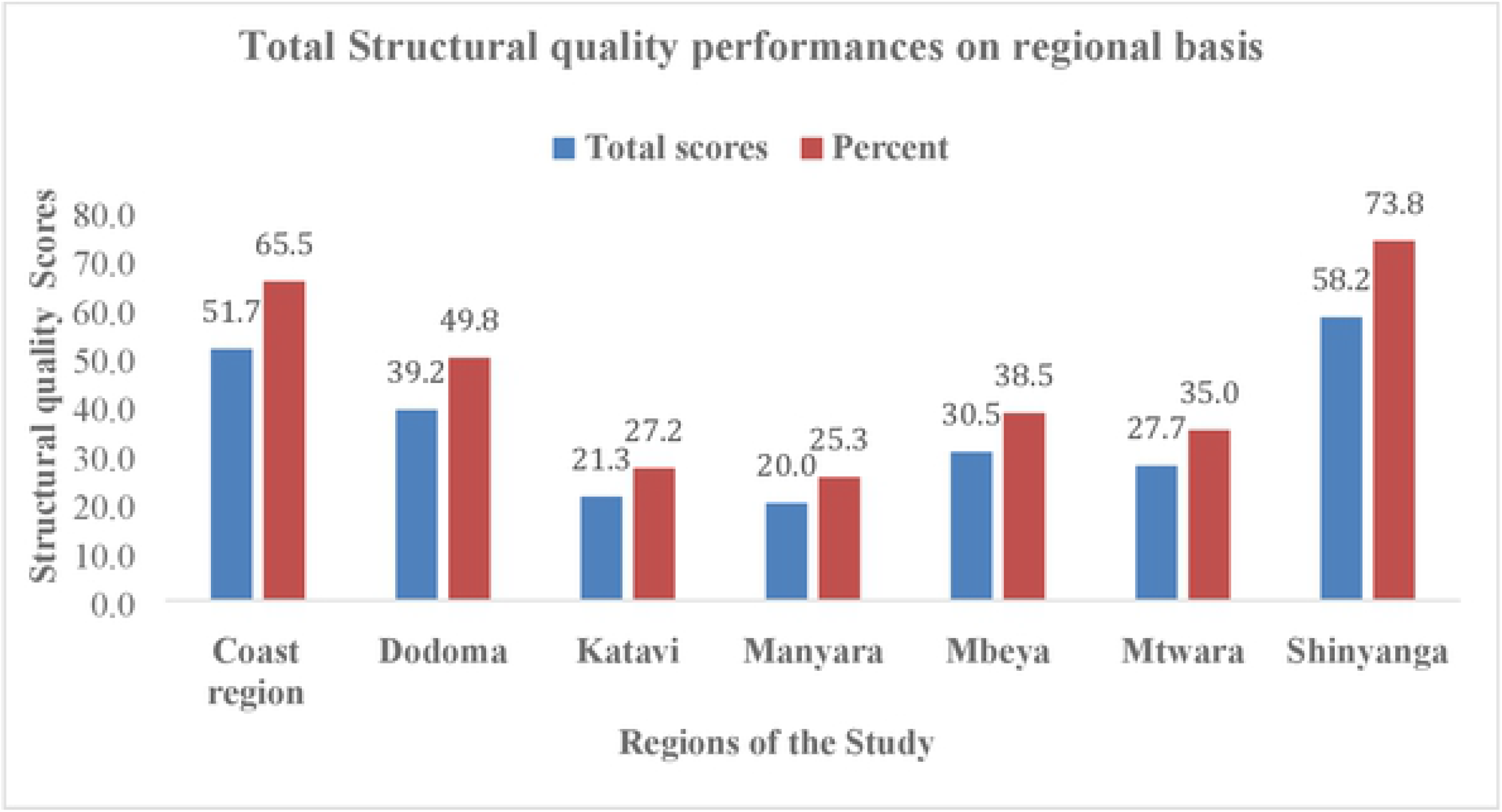
Total Structural Quality performances on regional basis

By comparing Dispensary and health centre performances on structural quality indicated relatively low differences among the attributes assessed. Specifically, they did not show statistical significant differences on privacy (p=1.000), hygiene and sanitation (p=.898), labour ward (p=.898), maternal health (p=.804) and waste management (p=.560). On the other hand, significant differences were observed on obstetric emergencies (p < .005), sterilization (p=.034) and overall structural quality (p=.018). Generally, health centre performed significantly higher on two attributes with obstetric emergencies indicating the highest magnitude of difference between dispensary and health centre (M=35.9, SD=20.3 and M=62.2, SD=20.9, respectively (Table 2).

With regard to rural-urban performance on structural quality, there was no statistical significant difference on total performance. Similarly, there was no significant differences between rural and urban health facilities on other assessed attributes of structural quality (p >.05) except for sterilization in which urban facilities performed significantly higher than the rural facilities [M=41.2, SD=27.7, 61.3, SD=28.4, respectively (p=. 028)]. On the other hand, marginal differences were observed on individual assessed attributes. For examples, rural facilities performed relatively higher than urban ones on privacy (41.2 and 32.0), maternal death (31.4 and 30.7) and waste management (49.0 and 47.3) respectively. Whereas, urban facilities performed relatively higher on all other assessed attributes (see the Table for clarifications). This means that, rural and urban health facilities could not differ significantly in terms of structural quality performance as an overall indicator (Table 3).

Multiple regression analysis was done to assess the relative effect of various explanatory variables or establish whether the predictor variables are independently associated with outcomes (dependent variable) of interest (Structural Quality attributes) was done to control/adjusting for any confounders as well. There is no single factor which shown to have been significant.

## Discussion

The objective of this study was to measure the structural quality of maternal health services at the primary health care facilities in Tanzania. The reason for undertaking this study is because maternal health remains to be the key priority of majority of Low Middle income countries including Tanzania. In this study, it was clear that majority of the regions had normal scores in terms of their structural quality in relation to the maternal health services except for two regions namely Shinyanga and Pwani which might have been due to the implementation of Results Based Financing (RBF) program. Structural quality indicators in the area around privacy, hygiene and obstetric emergences did score low as compared to other indicators, this is in line with another study which was done in which the said indicators also scored low [40]. Maternal death audit review remains a key challenge in most of primary health facilities in least developed countries as many deaths of mothers goes un checked which denies an opportunity to have lessons learnt from the deceased. From Donabedian model, the three domains namely; - structural, processes and outcome are important in defining the quality of any settings. Therefore high structural quality care processes remains to be a great challenge in this study and this could be reason which deters women from visiting health facilities for both antenatal and delivery services as also seen in another study done by Bishanga et al (2019) [41].

In this study, the structural quality comparison between Dispensary and Health center performances showed relatively low differences among the assessed attributes. Moreover, they did not show any statistical significant differences except for obstetric emergencies (p < .005), sterilization (p=. 034) and overall structural quality (p=. 018). The reason for statistical differences on the attribute of offering obstetric emergences might be due to fact that; most dispensaries do not have infrastructure to offer surgical services. With regard to rural-urban performance on structural quality, there was no statistical significant difference on total performance. Similarly, there was no significant differences between rural and urban health facilities on other assessed attributes of structural quality except for sterilization (p <. 05) and overall performance (p<0.028) in which urban facilities performed significantly higher than the rural facilities [M=41.2, SD=27.7, 61.3, SD=28.4]. This might be due to fact that there are more sterilization facilities in the urban as compared to the rural settings. Therefore, there is a need to improve the sterilization services in the rural settings, because if it is left unchecked may results into infections to the patients who receives surgical care services in those facilities. However some of the attributes the rural settings outperformed the urban settings, for examples, rural facilities performed relatively higher than urban ones on privacy (41.2 and 32.0), maternal death reviews (31.4 and 30.7) and waste management (49.0 and 47.3) respectively.

The above findings highlight the importance of surveys that will mainly focus from the process to the outcome instead of having surveys that mainly focuses on the inputs.

### Study Limitations

This study is limited, because it was a cross sectional study which is good to get snap shot findings however it does not give adequate evidence on the causal effect relationship. This study also excluded the private primary health facilities as they have varying degrees of service provision. Moreover, the data collection tool used was formulated basing on some previous studies and author’s experience.

## Conclusion

Generally facilities performed low on the structural quality indicators of maternal health services provision. However, they had high performance on two attributes that are sterilization and emergence obstetric care. Strengthening of Supportive supervision is needed so that to maintain good structural quality of maternal health services at the primary health care facility level.

## Abbreviations

CHMT: Council Health Management Team
DMO: District Medical Officer
DHIS-2: District Health Information System - 2
HFGC: Health Facility Governing Committee
HMIS: Health Management Information System
RBF: Results Based Financing
MoHCDGEC: Ministry of Health Community Development, Gender, Elderly and Children
PO-RALG: President’s Office – Regional Administration and Local Government
RAS: Regional Administrative Secretary
RHMT: Regional Health Management Team
PPHCF: Public Primary Health Care Facilities

## Authors’ contributions

NAK, AK, NHI, JB, and SMK developed and reviewed the Manuscript. All authors read and approved the final manuscript.

## Consent for Publication

All authors consented to publication.

---

Table 1: Descriptive statistics on type of health facilities performance on structural quality indicators

Table 2: Dispensary and health center performance on structural quality indicators

Table 3: Urban and Rural performance on structural quality indicators

**Additional File 1:**

## Reference

1. Mainz J, Bartels PD, Laustsen S et al. The National Indicator Project for monitoring and improving medical technical care [in Danish]. Ugeskr Laeger, 2001; 163: 6401–6406.

2. WHO (2001): Crossing the Quality Chasm: A New Health System for the 21st Century. Committee on Quality of Health Care in America, Institute of Medicine. Washington, DC, USA: National Academies Press; 2001

3. Donabedian A. Evaluating the quality of medical care. Milbank Mem Fund Q 1966;44(Suppl):166–206. 23.

4. Brand CA, Barker AL, Morello RT, et al. A review of hospital characteristics associated with improved performance. Int J Qual Health Care 2012; 24:483–94. 24.

5. Baker G, MacIntosh-Murray A, Porcellato C, Dionne L, Stelmacovich K, Born K. High Performing Healthcare Systems: Delivering Quality by Design. Toronto, Ontario, Canada: Longwoods Publishing Corporation, 2008.

6. Trends in Maternal Mortality: 1990 to 2015 Estimates by WHO, UNICEF, UNFPA, World Bank Group and the United Nations Population Division. It can be accessed at

7. https://apps.who.int/iris/bitstream/handle/10665/193994/WHO_RHR_15.23_eng.pdf;jsessionid=94661FFA0427719AEA0C8338D1FF282E?sequence=1

8. Kinney MV^1^, Kerber KJ, Black RE, Cohen B, Nkrumah F, Coovadia H, Nampala PM, Lawn JE; Science in Action: Saving the lives of Africa’s Mothers, Newborns, and Children working group, Axelson H, Bergh AM, Chopra M, Diab R, Friberg I, Odubanjo O, Walker N, Weissman E (2015): Sub-Saharan Africa’s mothers, newborns, and children: where and why do they die?. PLoS Med. 2010 Jun 21;7(6):e1000294. doi: 10.1371/journal.pmed.1000294. It can be accessed at https://www.ncbi.nlm.nih.gov/pubmed/20574524. It can be accessed at https://www.ncbi.nlm.nih.gov/pubmed/20574524

9. URT. 2016. Tanzania Demographic Health Survey and Malaria Indicator Survey (TDHS) 2015-2016. Ministry of Health Community Development, Gender, Elderly and Children. Ministry of Health Zanzibar. National Bureau of Statistics. Office of Chief Government Statistician Zanzibar.ICF. Rockville Maryland USA. https://www.nbs.go.tz/nbs/takwimu/dhs/2015-16_TDHS-MIS_Key_Findings_English.pdf. Accessed 13 Oct 2018.

10. MoHSW. (2015). Tanzania Health Sector Strategic Plan 2015-2020 (HSSP IV), Dar es Salaam. Pg. 53

11. MoHSW (2015): The National Road Map Strategic Plan to Improve Reproductive Health (One Plan II 2016-2020). It can be accessed https://www.globalfinancingfacility.org/sites/gff_new/files/Tanzania_One_Plan_II.pdf

12. National Bureau of Statistics (NBS) - Government of Tanzania, Office of Chief Government Statistician, Zanzibar (OCGS) - Government of Tanzania (2016): Tanzania - Service Provision Assessment Survey 2014–2015. It can be accessed at http://microdata.worldbank.org and file:///Users/dr.ntulikapologwe/Downloads/ddi-documentation-english_microdata-2573.pdf

13. Anne Austin, Ana Langer, Rehana A Salam, Zohra S Lassi, Jai K Das and Zulfiqar A Bhutta (2014): Approaches to improve the quality of maternal and newborn health care: an overview of the evidence. It can be accessed Reproductive Health 2014 11 (Suppl 2): S1 https://doi.org/10.1186/1742-4755-11-S2-S1. It can be accessed at https://reproductive-health-journal.biomedcentral.com/articles/10.1186/1742-4755-11-S2-S1

14. 1.Anatole, M., Magge, H., Redditt, V., Karamaga, A., Niyonzima, S., Drobac, P., Mukherjee, J.S., Ntaganira, J., Nyirazinyoye, L. and Hirschhorn, L.R. (2013), “Nurse mentorship to improve the quality of health care delivery in rural Rwanda”, Nursing Outlook, Vol. 61 No. 3, pp. 137–144. [Crossref], [Medline], [ISI], [Google Scholar] [Infotrieve]

15. Berman, J., Nkabane, E.L., Malope, S., Machai, S., Jack, B. and Bicknell, W. (2012), “Developing a hospital quality improvement initiative in Lesotho”, International Journal of Health Care Quality Assurance, Vol. 27 No. 1, pp. 15–24. [Link], [Google Scholar] [Infotrieve]

16. Das, J., Kumar, R., Salam, R., Lassi, Z. and Bhutta, Z. (2014), “Evidence from facility level inputs to improve quality of care for maternal and newborn health: interventions and findings”, Reproductive Health, Vol. 11 No. S2, p. 4. [Crossref], [Medline], [Google Scholar][Infotrieve]

17. Faye, A., Dumont, A., Ndiaye, P. and Fournier, P. (2014), “Development of an instrument to evaluate intrapartum care quality in Senegal: evaluation quality care”, International Journal for Quality in Health Care, Vol. 26 No. 2, pp. 184–189. [Crossref], [Medline], [ISI], [Google Scholar] [Infotrieve]

18. Ishijima, H., Eliakimu, E., Takahashi, S. and Miyamoto, N. (2014), “Factors influencing national rollout of quality improvement approaches to public hospitals in Tanzania”, Clinical Governance, Vol. 19 No. 2, pp. 137–152. [Link], [Google Scholar] [Infotrieve]

19. Dumont, A., Fournier, P., Abrahamowicz, M., Traoré, M., Haddad, S. and Fraser, W.D.(2013), “Quality of care, risk management, and technology in obstetrics to reduce hospital-based maternal mortality in Senegal and Mali (QUARITE): a cluster-randomised trial”, The Lancet, Vol. 382 No. 9887, pp. 146–157. [Crossref], [Medline], [ISI], [Google Scholar] [Infotrieve]

20. Kim, Y.M., Chilila, M., Shasulwe, H., Banda, J., Kanjipite, W., Sarkar, S., Bazant, E., Hiner, C., Tholandi, M., Reinhardt, S., Mulilo, J.C. and Kols, A. (2013), “Evaluation of a quality improvement intervention to prevent mother-to-child transmission of HIV (PMTCT) at Zambia defence force facilities”, BMC Health Services Research, Vol. 13 p. 345. [Crossref], [Medline], [ISI], [Google Scholar]

21. Colbourn, T., Nambiar, B., Bondo, A., Makwenda, C., Tsetekani, E., Makonda-Ridley, A., Msukwa, M., Barker, P., Kotagal, U., Williams, C., Davies, R., Webb, D., Flatman, D., Lewycka, S., Rosato, M., Kachale, F., Mwansambo, C. and Costello, A. (2013), “Effects of quality improvement in health facilities and community mobilization through women’s groups on maternal, neonatal and perinatal mortality in three districts of Malawi: MaiKhanda, a cluster randomized controlled effectiveness trial”, International Health, Vol. 5 No. 3, pp. 180–195. [Crossref], [Medline], [ISI], [Google Scholar] [Infotrieve]

22. Ali Mohammad Mosadeghrad (2014). *Factors influencing healthcare service quality. Int J Health Policy Manag. 2014 Jul; 3(2): 77–89. Published online 2014 Jul 26. doi: 10.15171/ijhpm.2014.65 PMCID: PMC4122083, PMID: 25114946

23. MoHSW (2011): Tanzania Quality Improvement Framework in Health care 2011-2016. Can be accessed at https://www.jica.go.jp/project/tanzania/006/materials/ku57pq00001x6jyl-att/framework_in_health.pdf

24. MoHSW (2013); NATIONAL HEALTH AND SOCIAL WELFARE QUALITY IMPROVEMENT STRATEGIC PLAN 2013 - 2018 (NHSWQISP-I - 2013 - 2018) NOVEMBER 2013. It can be accessed at http://www.tzdpg.or.tz/fileadmin/documents/dpg_internal/dpg_working_groups_clusters/cluster_2/health/Sub_Sector_Group/Quality_Assurance/11.b_NATIONAL_QI_STRATEGIC_PLAN_-_FINAL-isbn.pdf

25. Talhiya Yahya & Mohamed Mohamed (2018): Raising a mirror to quality of care in Tanzania: the five-star assessment. Published Online September 5, 2018 http://dx.doi.org/10.1016/S2214-109X (18) 30348–6

26. Tanzania Demographic Health Survey and Malaria Indicator Survey (TDHS) 2015-2016. It can be accessed at https://dhsprogram.com/pubs/pdf/FR321/FR321.pdf

27. WHO, UNICEF, UNFPA, World Bank Group and the United Nations Population Division. (2015). Trends in maternal mortality: 1990 to 2015, It can be accessed https://www.afro.who.int/sites/default/files/2017-05/trends-in-

28. Ntuli A. Kapologwe, Albino Kalolo, Stephen M. Kibusi, Zainab Chaula, Anna Nswilla, Thomas Teuscher, Kyaw Aung, Josephine Borghi. (2018). Understanding the implementation of Direct Health Facility Financing (DHFF) and its effect on health system performance in Tanzania: A non-controlled before and after mixed method study protocol. Journal: Health Research Policy and Systems. DOI: 10.1186/s12961-018-0400-3

29. Gilson L^1^, Magomi M, Mkangaa E (1995): The structural quality of Tanzanian primary health facilities. Bull World Health Organ. 1995; 73(1): 105–14. It can be accessed at https://www.ncbi.nlm.nih.gov/pubmed/7704920

30. Ministry of Health and Social Welfare (MoHSW) (2014): Results Based Financing Operational Manual. It can be accessed at https://bluesquarehub.files.wordpress.com/2017/03/tanzania-operations-manual-20160422_rbf_approved-version.pdf

31. Anne-Emanuelle Birn (2018). Back to Alma-Ata, From 1978 to 2018 and Beyond. Am J Public Health. 2018 September; 108(9): 1153–1155. Published online 2018 September. doi: 10.2105/AJPH.2018.304625. It an be accessed at https://www.ncbi.nlm.nih.gov/pmc/articles/PMC6085028/

32. The Astana Declaration: the future of primary health care? Published: The Lancent; October 20, 2018. Can be accessed at http://www.thelancet.com Vol 392 October 20, 2018. https://doi.org/10.1016/S0140-6736(18)32478-4

33. URT, Joint Ministry of Health and Social Welfare (MoHSW) and President’s Office – Regional Adimistration and Local government (PORALG) (2004): Health Basket and Health Block Grants Guidelines for the Disbursement of Funds, Preparation of Comprehensive Council Health Plans, Financial and Technical Reports and Rehabilitation of PHC Facilities by Councils. Can be accessed at http://ihi.eprints.org/804/1/MoHSW.pdf_%2831%29.pdf

34. Cohen, D. J., & Crabtree, B. F. (2008). Research in Health Care: Controversies and Recommendations. Annals Of Family Medicine, 6(4), 331–339. https://doi.org/10.1370/afm.818.INTRODUCTION

35. Ary, D., Jacobs, L., Sorenson, C., Razavieh, A., Jacobs, C., C, S., & Razavier, A. (2010). Introduction to Research in Education.

36. Kanji, Najmi & Kilima, Peter & Lorenz, Nicolaus & Garner, Paul. (1995). Quality of primary outpatient services in Dar-es-Salaam: A comparison of government and voluntary providers. Health policy and planning. 10. 186–90. 10.1093/heapol/10.2.186. It can be accessed at https://www.ncbi.nlm.nih.gov/pubmed/10143456

37. The World Bank. (2013). Results-Based Financing for Health. African Health Forum, 3–6.

38. Christoph Boller,1 Kaspar Wyss,2 Deo Mtasiwa,3 & Marcel Tanner.(2003). Quality and comparison of antenatal care in public and private providers in the United Republic of Tanzania. Bulletin of the World Health Organization 2003, 81 (2).

39. Ministry of Health and Social Welfare (MoHSW) (2014): Results Based Financing Operational Manual. It can be accessed at https://bluesquarehub.files.wordpress.com/2017/03/tanzania-operations-manual-20160422_rbf_approved-version.pdf

40. Mundodan JM. (2015). Quality of antenatal care services in selected health facilities of Kaski district, Nepal International Journal of Community Medicine and Public Health. 2015 Nov; 2(4): 513–519. It can be accessed at http://www.ijcmph.com [accessed Dec 28 2018]. DOI: 10.18203/2394-6040.ijcmph20182142

41. Hannah H. Leslie, Zeye Sun, Margaret E. Kruk (2017): Association between infrastructure and observed quality of care in 4 healthcare services: A cross-sectional study of 4,300 facilities in 8 countries. https://doi.org/10.1371/journal.pmed.1002464. It can be accessed at https://journals.plos.org/plosmedicine/article?id=10.1371/journal.pmed.1002464

42. Dunstan R. Bishanga, Joseph Massenga, Amasha H. Mwanamsangu,Young-Mi Kim, John George, Ntuli A. Kapologwe,Jeremie Zoungrana, Mary Rwegasira, Adrienne Kols, Kathleen Hill, Marcus J. Rijken and Jelle Stekelenburg (2019): Women’s experience of facility –based childbirth cares and receipt of an early postnatal checks for herself and her newborn in northwestern Tanzania. Int. J. Environ. Res. Public Health 2019, 16(3), 481; https://doi.org/10.3390/ijerph16030481

43. MoHSW (2014): STAFFING LEVELS FOR MINISTRY OF HEALTH AND SOCIAL WELFARE DEPARTMENTS, HEALTH SERVICE FACILITIES, HEALTH TRAINING INSTITUTIONS AND AGENCIES 2014-2019 (REVISED): It can be accessed at https://www.jica.go.jp/project/tanzania/006/materials/ku57pq00001x6jyl-att/REVIEW_STAFFING_LEVEL_2014-01.pdf

44. Tanzania Health Facility Registry (2018). It can be accessed at http://hfrportal.ehealth.go.tz

